# Enzymatic Beacons for Specific Sensing of Dilute Nucleic Acid and Potential Utility for SARS-CoV-2 Detection

**DOI:** 10.1101/2021.08.30.458287

**Authors:** Xiaoyu Zhang, Venubabu Kotikam, Eriks Rozners, Brian P. Callahan

## Abstract

Enzymatic beacons, or E-beacons, are 1:1 bioconjugates of the nanoluciferase enzyme linked covalently at its C-terminus to hairpin forming DNA oligonucleotides equipped with a dark quencher. We prepared E-beacons biocatalytically using the promiscuous “hedgehog” protein-cholesterol ligase, HhC. Instead of cholesterol, HhC attached nanoluciferase site-specifically to mono-sterylated hairpin DNA, prepared in high yield by solid phase synthesis. We tested three potential E-beacon dark quenchers: Iowa Black, Onyx-A, and dabcyl. Prototype E-beacon carrying each of those quenchers provided sequence-specific nucleic acid sensing through turn-on bioluminescence. For practical application, we prepared dabcyl-quenched E-beacons for potential use in detecting the COVID-19 virus, SARS-CoV-2. Targeting the E484 codon of the SARS-CoV-2 Spike protein, E-beacons (80 × 10^−12^ M) reported wild-type SARS-CoV-2 nucleic acid at ≥1 × 10^−9^ M with increased bioluminescence of 8-fold. E-beacon prepared for the E484K variant of SARS-CoV-2 functioned with similar sensitivity. These E-beacons could discriminate their complementary target from nucleic acid encoding the E484Q mutation of the SARS-CoV-2 Kappa variant. Along with specificity, detection sensitivity with E-beacons is two to three orders of magnitude better than synthetic molecular beacons, rivaling the most sensitive nucleic acid detection agents reported to date.

---

Reagents for the specific detection of dilute nucleic acid are fundamental to molecular and cellular genetics and various clinical diagnostics, including tests for viral pathogens like SARS-CoV-2.*^1–4^* Molecular beacons, the fluorogenic hairpin-forming oligonucleotide-based sensors, have provided a standard detection tool.*^5, 6^* Modified with fluorophore and quencher at opposite ends, the oligonucleotide fluorescence is suppressed in the hairpin, or off state. Molecular beacons switch on by hybridizing to complementary nucleic acid, which separates the fluorophore and quencher, increasing radiative emission. Improvements to molecular beacon technology have been driven mainly through chemical synthesis with the introduction of more efficient quenchers and new fluorophore/quencher pairs.*^7^*

Here we describe an alternative strategy where enzymatic catalysis is employed in the preparation of the sensor and for the sensor output. In enzymatic beacons or E-beacons the fluorophore of a molecular beacon is replaced by the compact, ATP-independent bioluminescent enzyme, nanoluciferase (Nluc).*^8–11^* This substitution provides an internal, amplifiable light source. Nluc with the engineered substrate furimazine produces light that is sufficiently bright for measurement with portable luminometer.*^12^* Moreover, we expected that nucleic acid detection assays would consume less reagent while improving detection sensitivity relative to synthetic molecular beacons.

To connect Nluc to a hairpin-forming oligonucleotide as a potential E-beacon, we applied the protein-nucleic acid bioconjugation activity of hedgehog proteins **(Figure 1A)**.*^13^* Hedgehog proteins harbor a promiscuous autoprocessing domain, HhC, that catalyzes two linked activities: self-cleavage from an adjacent N-terminal protein, and site-specific sterol ligation to that departing N-terminal protein. These reactions occur simultaneously without cofactor requirements. We have found that the *Drosophila melanogaster* HhC has broad substrate tolerance, maintaining robust bioconjugation activity toward heterologous substrate proteins and catalyzing ligation with sterols of varying structure and chemical appendages, including oligonucleotides. *^13, 14^* For E-beacons, we fused this promiscuous HhC to the C-terminus of the Nluc enzyme **(Supporting Methods).**

**FIGURE 1.**
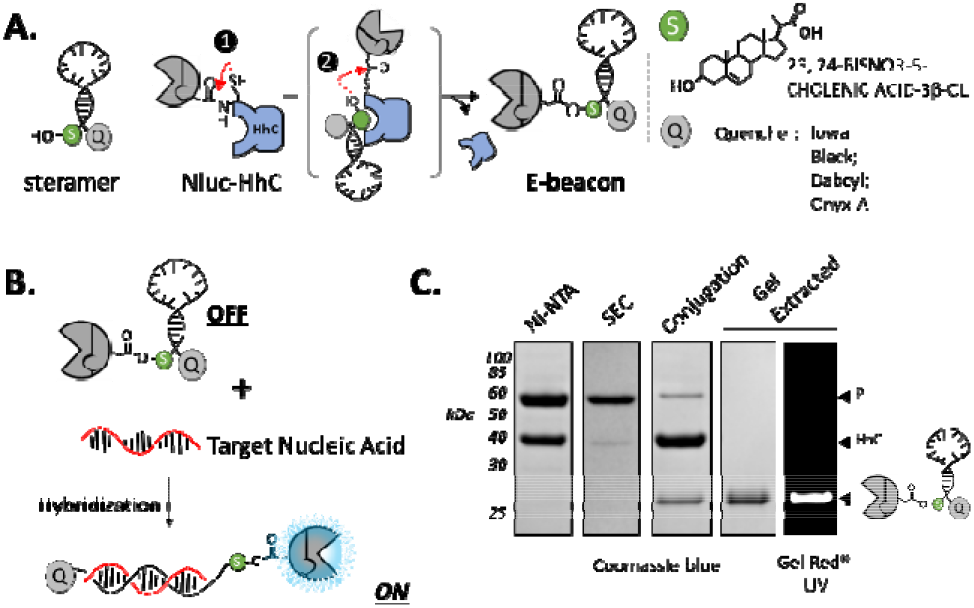
Concept and biocatalytic preparation of prototype E-beacon (A) Scheme showing HhC catalyzed bioconjugation of Nluc (grey) to a hairpin-forming mono-sterylated oligonucleotide (steramer) equipped with a 3’ dark quencher (Q). Conjugation involves formation of an internal thioester (step 1), followed by steramer binding and acyl transfer to the sterol hydroxyl group (step 2). (B) General mechanism for turn-on nucleic acid detection by E-beacon, whereby hybridization of hairpin oligonucleotide with complementary nucleic acid displaces the quencher (Q) from the Nluc enzyme, enhancing bioluminescence signal. (C) E-beacon preparation monitored by SDS-PAGE. A fusion of Nluc with HhC is purified first by Ni-NTA chromatography, followed by size exclusion chromatography (SEC), then reacted with steramer (conjugation). E-beacon is isolated by agarose gel extraction. Nucleic acid was visualized by UV with GelRed staining.

To serve as a substrate for ligation to Nluc, the hairpin oligonucleotide component required mono-sterylation. Our initial tests used an ssDNA oligo with the sequence: (5’-amino)-<u>CGCTC</u>*CCAAAAAAAAAAACC*<u>GAGCG</u>-(3’-IBQ). The underlined regions self-anneal to form the hairpin stem; the italicized 15 nucleotides in the middle represent the probe region for target nucleic acid hybridization. The same sequence was used by Kramer, Tyagi et al for their early biophysical studies on molecular beacons.*^15^* We used the IowaBlack (IBQ) dark quencher at the 3’-end to suppress Nluc bioluminescence in the hairpin (off) state **(Figure 1B).** Mono-sterylation was achieved by coupling the oligo’s 5’-amino group to the carboxyl group of 23,24-bisnor 5-cholenic acid **(Supporting Methods)**.^a^

We prepared a prototype E-beacon, Eb.1, by combining the Nluc-HhC precursor protein with mono-sterylated hairpin-forming oligonucleotide. His-tagged Nluc-HhC precursor was expressed in *E. coli* and purified in soluble form by Ni-NTA resin and size exclusion chromatography **(Figure 1C**, lanes 1-2). Site-specific bioconjugation of Nluc to the modified hairpin was carried out in vitro.*^13^* In a typical reaction, we combined Nluc-HhC precursor at 2 μM (final) with 25-50-fold excess of hairpin in HhC buffer, incubated on the benchtop overnight, and assessed reaction progress by SDS-PAGE **(Figure 1C**, see conjugation). Agarose gel extraction provided a convenient means of purifying the conjugate, Eb.1 **(Figure 1C**, last two lanes).

We observed encouraging nucleic acid sensing with Eb.1. As an initial test, bioluminescence readings from samples with Eb.1 plus oligonucleotide complementary to the hairpin probe region (signal; GGTTTTTTTTTTTGG) were collected and compared with bioluminescence readings from Eb.1 mixed with non-complementary oligonucleotide (noise; CTGGTCTTCGGGCTA). Eb.1 was present at 2 × 10^−9^ M, oligonucleotide was added to 25 × 10^−9^ M. Samples were incubated in hybridization buffer (KCl 100 mM, MgCl_2_ 1 mM, in 10 mM Tris buffer, pH 8) at 25 °C for 30 min. The Nluc substrate furimazine was then added according to manufacturer instructions (Promega).*^8^* Representative data from a 96-well experiment are summarized in **Figure 2A.** The average ratio of Eb.1 signal (with target) to noise (with random oligo) was 2.5-3-fold. The increased bioluminescence from Eb.1 in samples with complementary oligo accords with the unquenching mechanism*^5, 6^*: target/probe hybridization displaces the 3’quencher from the light source, here Nluc, permitting brighter signal.^b^

**FIGURE 2.**
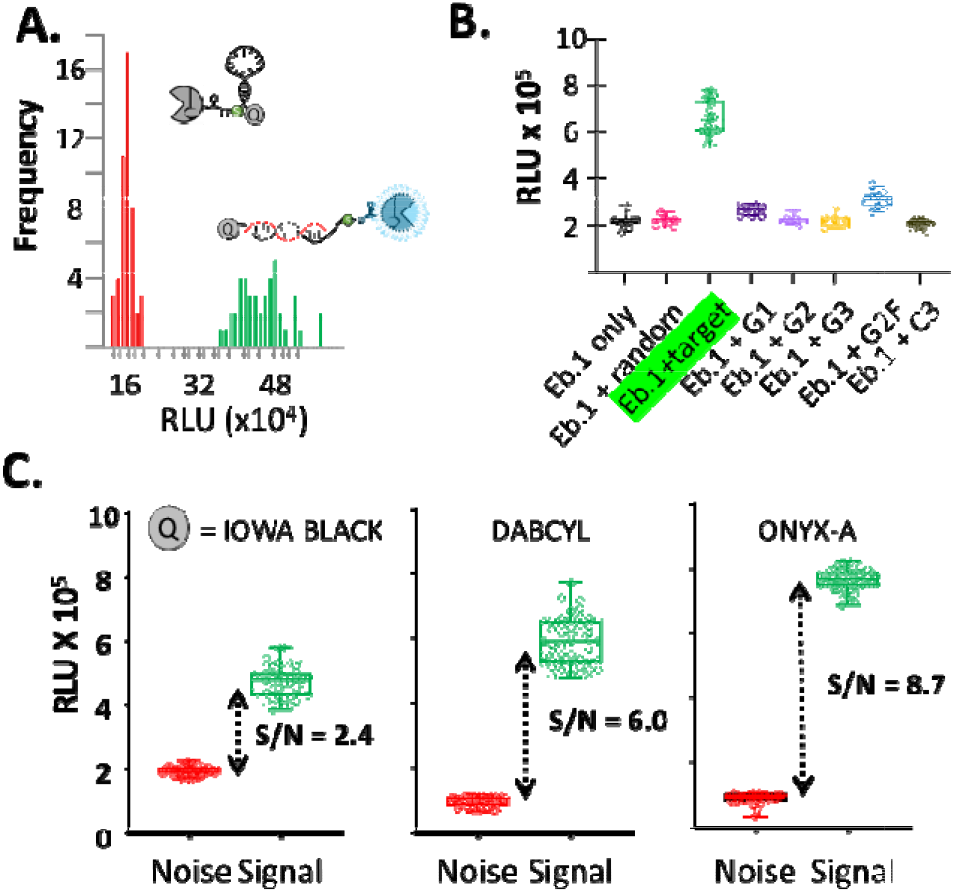
E-beacon proof of concept and sequence specific nucleic acid detection*. **(A)** Bioluminescence from Eb.1 increases in the presence of complementary oligonucleotide (green, n=48) compared to E-beacon samples mixed with noncomplementary oligonucleotide (red, n=48). **(B)** Results of “blinded test” of Eb.1 specificity indicate that Eb.1 distinguishes complementary oligonucleotide (green) from oligonucleotides containing 1-3 base mismatches. See Table 1 for sequence information. **(C)** Comparison of E-beacons with different quenchers: IowaBlack, dabcyl, and ONYX-A. Bioluminescence was measured after 10-minute incubation with complementary oligonucleotide (*green*, n=48) or noncomplementary oligonucleotide (*red*, n=48). *In **A, B**, and **C**, the E. beacon was present at 2 × 10^−9^ M final; oligonucleotide at 25 × 10^−9^ M; temperature, 25 °C; substrate, furimazine.

Next, we more stringently evaluated the specificity of Eb.1 as a nucleic acid reporter by measuring bioluminescence output when combined with oligonucleotides carrying only one to three mismatches with the hairpin probe region. Oligonucleotide sequences are listed in **Table 1.** As above, each oligonucleotide was added to 25 × 10^−9^ M in hybridization buffer containing Eb.1 at 2 × 10^−9^ M. The experiments were carried out “blinded”: one member of the lab distributed complementary and mismatched oligonucleotides into sample wells of a 96-well plate in a semi-random pattern; another lab member without knowledge of the plate layout added Eb.1 and carried out the bioluminescence measurement and data analysis. The results of a representative specificity test are summarized in **Figure 2B.** The largest difference in Eb.1 bioluminescence is apparent between the perfectly complementary target and the fully mismatched template (signal/noise, 3-fold). No increase in bioluminescence was apparent in wells containing oligo with mismatches of two and three base pairs (G2 and G3). Oligo with one mismatch (G1) in the center of the probe sequence produced 10% unquenching of the E-beacon. Unquenching of 15-20% of Eb.1 was observed when mixed with oligo G2F, carrying (T-to-G) mismatches to the probe region at both the 3’ and the 5’ flanks. Nonetheless, samples of Eb.1 with G1 and G2F, compared to samples with Eb.1 plus the complementary oligo were easily distinguished. The fidelity of Eb.1 observed here suggested that this prototype’s specificity, like a molecular beacon,*^5, 6^* was sufficiently strong for detecting subtle sequence variations.

**TABLE 1.**
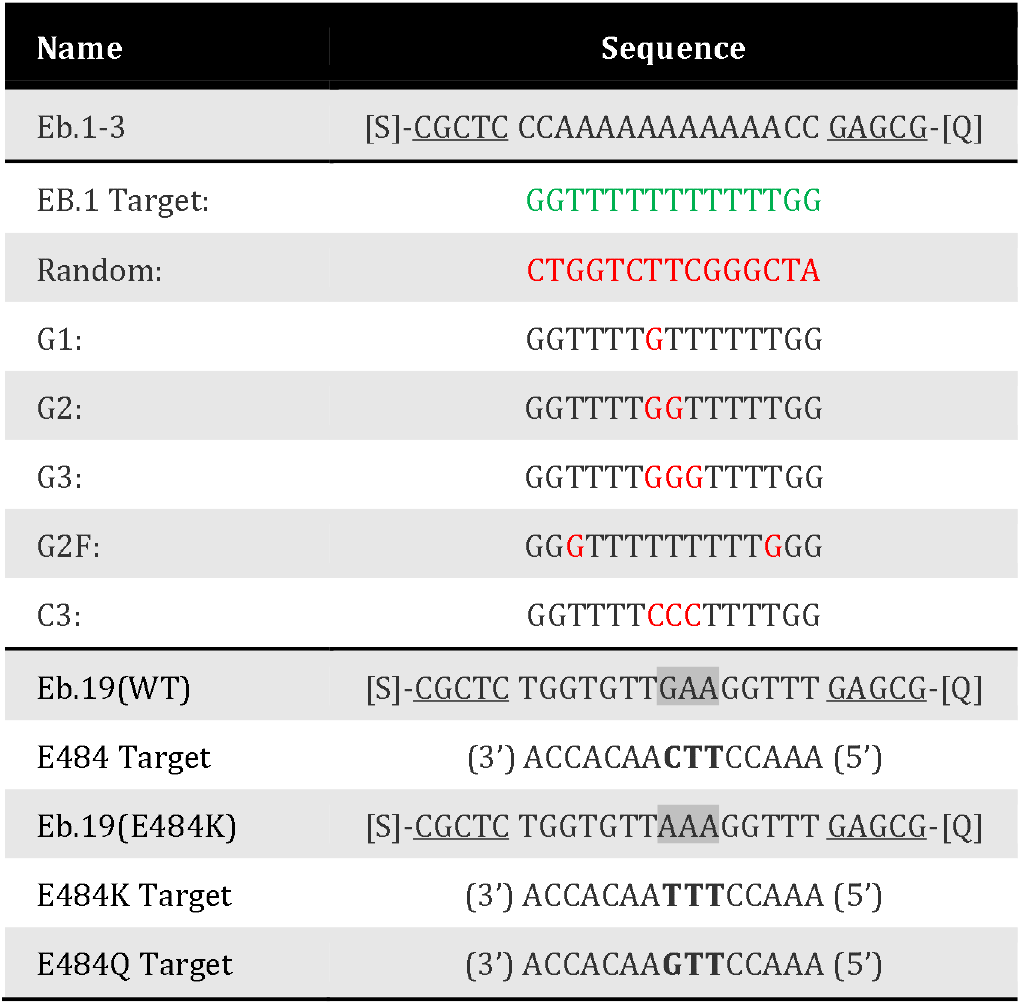
Sequences of hairpin oligonucleotides for Ebeacons and ssDNA oligonucleotides used for evaluating Ebeacon function. Red =mismatch; Bold = codon 484 of SARS-CoV2-2 spike protein; [S] sterol; [Q] quencher

While encouraged by the results above, we viewed the signal/noise range of ~3 as narrow (Z’ factor *^16^*, 0.46) and this led us to explore other quencher groups. The peak bioluminescence from the Nluc/furimazine reaction occurs at 460 nm. The maximum absorbance for the Iowa Black quencher is 531 nm. As alternative dark quenchers, we evaluated dabcyl and ONYX-A (Sigma). The dabcyl max absorbance peak is 453 nm and the ONYX_A absorbance peak is 515 nm. For meaningful comparison with Eb.1, we prepared new E-beacons from mono-sterylated oligonucleotide carrying the same sequence as Eb.1 and we applied the same biocatalytic approach for site-specific attachment to the C-terminus of Nluc. The dabcyl containing E-beacon, Eb.2, and the ONYX E-beacon, Eb.3, were likewise isolated by agarose gel extraction.

Changing the E-beacon quencher improved signal/noise and assay quality, in the order ONYX-A> dabcyl> IBQ. **Figure 2C** compares Eb.1 Eb.2 and Eb.3 readings with Ebeacons at 2 × 10^−9^ M and complementary or random oligo at 25 × 10^−9^ M. Eb.3 with the ONYX quencher showed the highest single/noise of 8.7-fold and Z’ of 0.79. Using the dabcyl quencher in Eb.2, the signal/noise was 6, better than IBQ, however the scatter in readings dropped the Z’ to 0.47. Because the oligonucleotide sequences in EB.1-.3 were the same, the changes in signal/noise are difficult to explain in terms of intra-oligo interactions, solution properties of the E-beacons, or Nluc activity; they seem more likely the result of specific photochemical characteristics (e.g. quenching efficiency). Structural information on SIGMA’s ONYX quencher which might have helped interpret these results is not yet available.

Lastly and with a view toward application, we designed and tested prototype E-beacons for SARS-CoV-2 detection. Reliable assays for rapid identification of patients infected with SARS-CoV-2, particularly those shedding viable virus, remain crucial to pandemic management. In considering how to configure the E-beacon for high-volume testing, we settled on using the dabcyl quencher because it is widely available. We also shifted our oligonucleotide sterylation protocol from microscale EDC-based coupling to a more robust solid phase method with a DNA synthesizer. The solid phase synthesis routinely provided >90% oligonucleotide sterylation efficiency and the modified oligos could be used without additional purification for Nluc conjugation. We found that E-beacon prepared this way displayed favorable S/N of 8-9 fold and Z’ factor >0.8 in test assays (see Supporting Figure 1 and Supporting Methods).

We selected the spike protein of SARS-CoV-2 as target for this new set of E-beacons. We focused on the region encompassing glutamate 484 of spike as changes at this codon can distinguish wild-type SARS-CoV-2 from emerging variants. The E-beacon intended for wild-type, Eb.19(WT), was prepared by HhC catalyzed ligation of Nluc to the sterylated hairpin: (sterol)-5’-<u>CGCTC</u>TGGTGTT**GAA**GGTTT<u>GAGCG</u>-3’-(dabcyl). A second E-beacon was prepared for detecting the E484K mutation, which is present in certain α and all β and γ strains of SARS-CoV-2. This Eb.19(E484K) carried the hairpin oligo: (sterol)-5’-<u>CGCTC</u>TGGTGTT**AAA**GGTTT<u>GAGCG</u>-3’-(dabcyl). The bold text represents the spike 484 codon.

We explored potential assay conditions for the SARS Ebeacons in two stages. First, we sought to determine the minimum working concentration of E-beacon. In our experiments, the concentration of target oligonucleotide was held constant and in excess at 1 × 10^−7^ M, while the Ebeacon was titrated from 1 × 10^−8^ M to 1 × 10^−16^ M. Bioluminescence readings from substrate furimazine were recorded as before. A positive result was defined arbitrarily as having a signal/noise of >5, anything less was deeme dam biguous. With the S/N threshold at 5, the lowest possible operating concentration of Eb.19 (WT) was 3 × 10^−13^ M (Supporting Figure 2)^c^.

In stage two, we held the E-beacon concentration constant and titrated the test oligos to find the nucleic acid detection threshold. We chose to use 8 × 10^−11^ M for the Ebeacon. Although higher than the minimum Eb.19 (WT) concentration defined above, we selected 80 pM to avoid pitfalls associated with assaying biomolecules in the extremely dilute regime, such as slow association kinetics and idiosyncratic adsorption effects, i.e., “death by dilution”. The assay parameter we sought was the EC_50_ or the concentration of target oligo that could produce 50% unquenching of Eb.19 (WT). We also explored the effect of E-beacon/oligo incubation time on sensor output. In **Figure 3A**, we show bioluminescence readings of samples with Eb.19 (WT) plotted as a function of increasing target oligo concentration after 10 min incubation and 3 hr incubation. Hyperbolic binding isotherms could be fit to the data to derive EC_50_ values. For Eb.19 (WT) the EC_50_ was 1.64 × 10^−9^ M for the 10 min incubation; for 3 hr incubation of Eb.19(WT) and oligo, the assay was 15-fold more sensitive, EC_50_ of 0.11 × 10^−9^ M. Similar behavior was observed with Eb.19 (E484K) (Supporting Figure 3). The improved sensitivity (lower EC_50_) with a 3 hr incubation of Eb19 (WT) and target oligo indicates that “slow” probe/target hybridization should be considered when attempting to detect very dilute nucleic acid.

**FIGURE 3.**
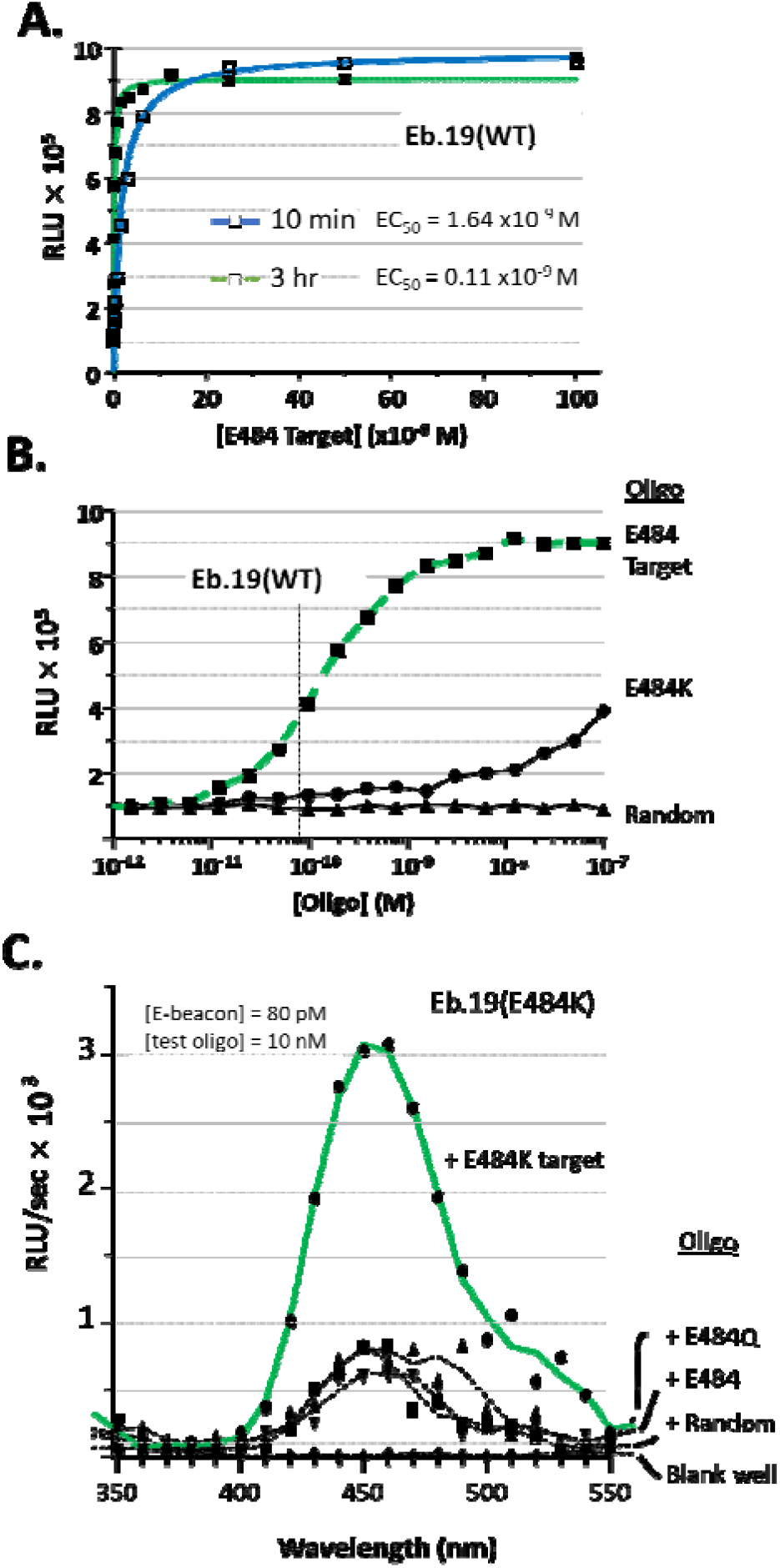
E-beacons for SARS-CoV-2 (A) Bioluminescence readings of samples with Eb. 19 (WT) after 10 min or 3 hr incubation with increasing concentration of complementary oligonucleotide. The curves show hyperbolic binding isotherms with calculated half maximum emission (EC_50_) at 2.54 × 10^−9^ M of oligonucleotide using the 10 min incubation, and 0.11 X 10^−9^ M for the 3 incubation. (B) Single base-pair mismatch discrimination by E-beacon, Eb. 19 (WT). Bioluminescence from Eb.19(WT), at 80 × 10^−12^ M, is plotted as a function of increasing oligonucleotide concentration. E484 is the complementary target, E484K is single base pair mismatch, “random” is non-complementary (see Table 1). (C) Bioluminescence spectrum showing the selective detection of E484K oligonucleotide by E-beacon, Eb.19 (E484K). Oligonucleotides E484 and E484Q represent single base pair variants of wild type SARS-CoV-2 and Kappa SARS-CoV-2, respectively.

To assess specificity, we compared signal from samples of Eb.19 (WT) mixed with the target oligo to signal from samples of Eb.19 (WT) mixed with the random oligo and the E484K variant oligo (single base-pair mismatch). As with the Eb.1 prototype, the random oligo did not unquench Eb. 19 (WT) over the entire concentration range tested (10^−12^ to 10^−7^ M) **(Figure 3B**, lowest trace). Unquenching of Eb. 19 (WT) did become apparent in samples containing the E484K oligo when added >10^−8^ M, although bioluminescence remained below samples with the fully complementary target oligo **(Figure 3B**, compare squares and circles). In **Figure 3C**, we present selectivity data for Eb.19 (E484K) in the form of comparative bioluminescence spectra. Here Eb.19 (E484K) was mixed with either E484K target oligo **(Figure 3C**, top trace), with random oligonucleotide, E484 (WT) or an oligo corresponding to E484Q of the SARS-CoV-2 Kappa variant. Once again, although some unquenching is apparent with base variants, signal of Eb. 19 (E484K) was greatest with the fully complementary target, E484K (6-8 fold). Together, the results in **Figure 3 A-C** are promising and show that with 80 pM E-beacon, nucleic acid detection is sensitive and base pair specific.

In summary, we report the preparation and analysis of E-beacons, enzymatic bioluminescent counterparts to synthetic fluorogenic molecular beacons. The general concept was introduced by Deo and her associates who assembled a (Renilla) luciferase-hairpin oligonucleotide-(3’-dabcyl) conjugate for microRNA detection,*^10^* and subsequently created a (Gaussia) luciferase-hairpin oligonucleotide-(3’-dabcyl) conjugate for HIV detection *^9^* and see also Ref *^17^*. The Deo sensors were prepared by spontaneous coupling of hairpin-forming oligos onto reactive residues of those luciferase enzymes. Spontaneous coupling is relatively inexpensive, but the technique produces heterogeneity in enzyme: oligonucleotide stoichiometry (1:1, 1:2, 1:3 etc) and the site(s) of attachment can vary between batches. We prepare E-beacons by site-specific C-terminal coupling of nanoluciferase using HhC as the bioconjugation catalyst.*^13^* This approach yields 1:1 enzyme: oligonucleotide stoichiometry. HhC catalyzed bioconjugation requires mono-sterylated oligonucleotides (steramers). Although steramers are not yet available commercially, we show here that they can be obtained in >90% yield from commercial reagents on a DNA synthesizer. E-beacons prepared by this approach for SARS-Cov-2 showed sensitive and sequence specific nucleic acid detection. Synthetic molecular beacons are used at ~ 1 × 10^−7^ M; the Deo sensors were used at 2 × 10^−9^ M; E-beacons with the ultrabright Nluc/furimazine reaction are effective in the mid to low pM range. Because such a small quantity of Ebeacon is needed for each assay, a micro-scale E-beacon preparation yields sufficient reagent for testing thousands of samples.

## ASSOCIATED CONTENT

### Supporting Information

(Word Style “Section_Content”). A listing of the contents of each file supplied as Supporting Information should be included. For instructions on what should be included in the Supporting Information as well as how to prepare this material for publication, refer to the journal’s Instructions for Authors.

The Supporting Information is available free of charge on the ACS Publications website. brief description (file type, i.e., PDF) brief description (file type, i.e., PDF)

## AUTHOR INFORMATION

### Notes

The Research Foundation of SUNY holds a patent for the general methodology used to prepare E-beacons. BPC is an inventor on this patent.

## ACKNOWLEDGMENT

We acknowledge generous support from the National Institute of Allergy and Infectious Diseases (Grant R03 AI163907 to B.P.C.), and the National Cancer Institute (Grant R01 CA206592 to B.P.C.), and the National Institute of General Medical Sciences (Grant R35 GM130207 to E.R.). We thank Callahan lab members Zihan Xu and Andrew Wagner for help with the “blinded” experiments and Daniel Powell for Nluc plasmids. We thank Dr. Juergen Schulte for expert help with NMR.

## Footnotes

a Oligonucleotide sterylation for the prototype E-beacons, Eb. 1-3, was achieved by EDC coupling. After butanol extraction and RP-HPLC purification, the final isolated yield of the sterylated oligo varied between 30-40%. The efficiency of oligonucleotide sterylation was improved significantly by switching to solid phase methods with a DNA synthesizer, *vide infra*.

b We compared hybridization driven unquenching of with nuclease driven unquenching. Digestion of the hairpin component with DNASE-1 produced similar bioluminescence enhancement compared hybridization driven unquenching (Supporting Figure 4).

c E-beacon measurements at this low concentration required setting the Biotek luminometer *gain* to the maximum level.

## REFERENCES

[1] Corman, V. M., Landt, O., Kaiser, M., Molenkamp, R., Meijer, A., Chu, D. K., Bleicker, T., Brunink, S., Schneider, J., Schmidt, M. L., Mulders, D. G., Haagmans, B. L., van der Veer, B., van den Brink, S., Wijsman, L., Goderski, G., Romette, J. L., Ellis, J., Zambon, M., Peiris, M., Goossens, H., Reusken, C., Koopmans, M. P., and Drosten, C. (2020) Detection of 2019 novel coronavirus (2019-nCoV) by real-time RT-PCR, Euro Surveill 25.

[2] Arons, M. M., Hatfield, K. M., Reddy, S. C., Kimball, A., James, A., Jacobs, J. R., Taylor, J., Spicer, K., Bardossy, A. C., Oakley, L. P., Tanwar, S., Dyal, J. W., Harney, J., Chisty, Z., Bell, J. M., Methner, M., Paul, P., Carlson, C. M., McLaughlin, H. P., Thornburg, N., Tong, S., Tamin, A., Tao, Y., Uehara, A., Harcourt, J., Clark, S., Brostrom-Smith, C., Page, L. C., Kay, M., Lewis, J., Montgomery, P., Stone, N. D., Clark, T. A., Honein, M. A., Duchin, J. S., Jernigan, J. A., Public, H.-S., King, C., and Team, C. C.-I. (2020) Presymptomatic SARS-CoV-2 Infections and Transmission in a Skilled Nursing Facility, N Engl J Med 382, 2081–2090.

[3] Hosseini, A., Pandey, R., Osman, E., Victorious, A., Li, F., Didar, T., and Soleymani, L. (2020) Roadmap to the Bioanalytical Testing of COVID-19: From Sample Collection to Disease Surveillance, ACS Sens 5, 3328–3345.

[4] Larremore, D. B., Wilder, B., Lester, E., Shehata, S., Burke, J. M., Hay, J. A., Tambe, M., Mina, M. J., and Parker, R. (2020) Test sensitivity is secondary to frequency and turnaround time for COVID-19 screening, Sci Adv.

[5] Marras, S. A., Kramer, F. R., and Tyagi, S. (1999) Multiplex detection of single-nucleotide variations using molecular beacons, Genet Anal 14, 151–156.

[6] Tyagi, S., and Kramer, F. R. (1996) Molecular beacons: probes that fluoresce upon hybridization, Nat Biotechnol 14, 303–308.

[7] Moutsiopoulou, A., Broyles, D., Dikici, E., Daunert, S., and Deo, S. K. (2019) Molecular Aptamer Beacons and Their Applications in Sensing, Imaging, and Diagnostics, Small 15, e1902248.

[8] Hall, M. P., Unch, J., Binkowski, B. F., Valley, M. P., Butler, B. L., Wood, M. G., Otto, P., Zimmerman, K., Vidugiris, G., Machleidt, T., Robers, M. B., Benink, H. A., Eggers, C. T., Slater, M. R., Meisenheimer, P. L., Klaubert, D. H., Fan, F., Encell, L. P., and Wood, K. V. (2012) Engineered luciferase reporter from a deep sea shrimp utilizing a novel imidazopyrazinone substrate, ACS Chem Biol 7, 1848–1857.

[9] Joda, H., Moutsiopoulou, A., Stone, G., Daunert, S., and Deo, S. (2018) Design of Gaussia luciferase-based bioluminescent stem-loop probe for sensitive detection of HIV-1 nucleic acids, Analyst 143, 3374–3381.

[10] Hunt, E. A., and Deo, S. K. (2011) Bioluminescent stem-loop probes for highly sensitive nucleic acid detection, Chem Commun (Camb) 47, 9393–9395.

[11] Biewenga, L., Rosier, B., and Merkx, M. (2020) Engineering with NanoLuc: a playground for the development of bioluminescent protein switches and sensors, Biochem Soc Trans 48, 2643–2655.

[12] Elledge, S. K., Zhou, X. X., Byrnes, J. R., Martinko, A. J., Lui, I., Pance, K., Lim, S. A., Glasgow, J. E., Glasgow, A. A., Turcios, K., Iyer, N. S., Torres, L., Peluso, M. J., Henrich, T. J., Wang, T. T., Tato, C. M., Leung, K. K., Greenhouse, B., and Wells, J. A. (2021) Engineering luminescent biosensors for point-of-care SARS-CoV-2 antibody detection, Nat Biotechnol.

[13] Zhang, X., Xu, Z., Moumin, D. S., Ciulla, D. A., Owen, T. S., Mancusi, R. A., Giner, J. L., Wang, C., and Callahan, B. P. (2019) Protein-Nucleic Acid Conjugation with Sterol Linkers Using Hedgehog Autoprocessing, Bioconjug Chem 30, 2799–2804.

[14] Ciulla, D. A., Wagner, A. G., Liu, X., Cooper, C. L., Jorgensen, M. T., Wang, C., Goyal, P., Banavali, N. K., Pezzullo, J. L., Giner, J. L., and Callahan, B. P. (2019) Sterol A-ring plasticity in hedgehog protein cholesterolysis supports a primitive substrate selectivity mechanism, Chem Commun (Camb) 55, 1829–1832.

[15] Bonnet, G., Tyagi, S., Libchaber, A., and Kramer, F. R. (1999) Thermodynamic basis of the enhanced specificity of structured DNA probes, Proc Natl Acad Sci U S A 96, 6171–6176.

[16] Zhang, J. H., Chung, T. D., and Oldenburg, K. R. (1999) A Simple Statistical Parameter for Use in Evaluation and Validation of High Throughput Screening Assays, J Biomol Screen 4, 67–73.

[17] Engelen, W., van de Wiel, K. M., Meijer, L. H. H., Saha, B., and Merkx, M. (2017) Nucleic acid detection using BRET-beacons based on bioluminescent protein-DNA hybrids, Chem Commun (Camb) 53, 2862–2865.

