## Supplemental Information for "Enzymatic Beacons for Specific Sensing of Dilute Nucleic Acid and Potential Utility for SARS-CoV-2 Detection"

### CONTENTS

Supporting Figures 1-4 ………….. 2-3

Materials…………………………. 4

Protein Expression/Purification….. 5

EDC-based steramer synthesis….. 6

Sterol amine synthesis…………… 7

Solid Phase Steramer Synthesis…… 9

E-beacon assembly by HhC……… 11

E-beacon isolation…………………. 11

E-beacon bioluminescence……….. 12

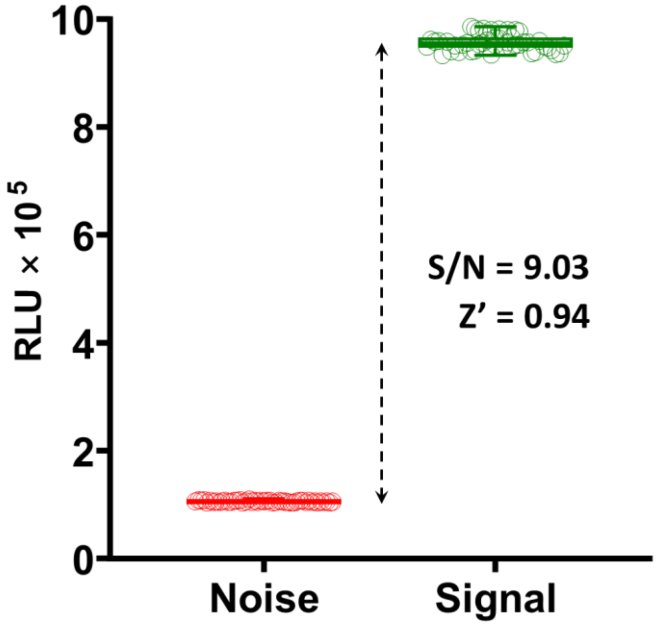


**Supporting Figure 1.** E-beacon (Eb.2') prepared with solid phase synthesized mono-sterylated oligo displays favorable signal to noise ratio and limited data scatter. Bioluminescence from samples with Eb.2' was measured after 10-minute incubation with complementary oligonucleotide (Signal, green, n=48) or with noncomplementary oligonucleotide (Noise, red, n=48). The E-beacon was present at $8 \times 10^{-11}$ M final; oligonucleotide at $1 \times 10^{-7}$ M; temperature, 25 °C; substrate, furimazine.

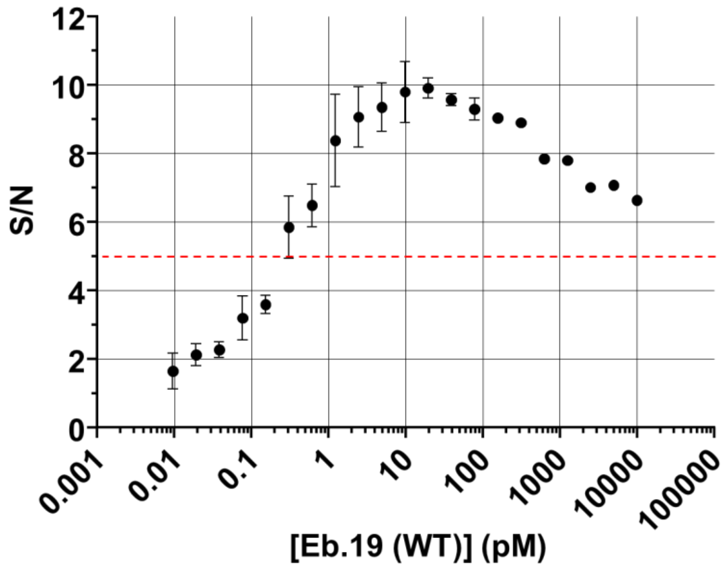


**Supporting Figure 2.** S/N ratio exceeds 5-fold with sub-picomolar Eb.19 (WT). Target oligonucleotides was held constant at $1 \times 10^{-7}$ M while the Eb.19 (WT) was titrated from $1 \times 10^{-8}$ M to $1 \times 10^{-16}$ M. Signal to noise ratios are plotted on the Y axis.

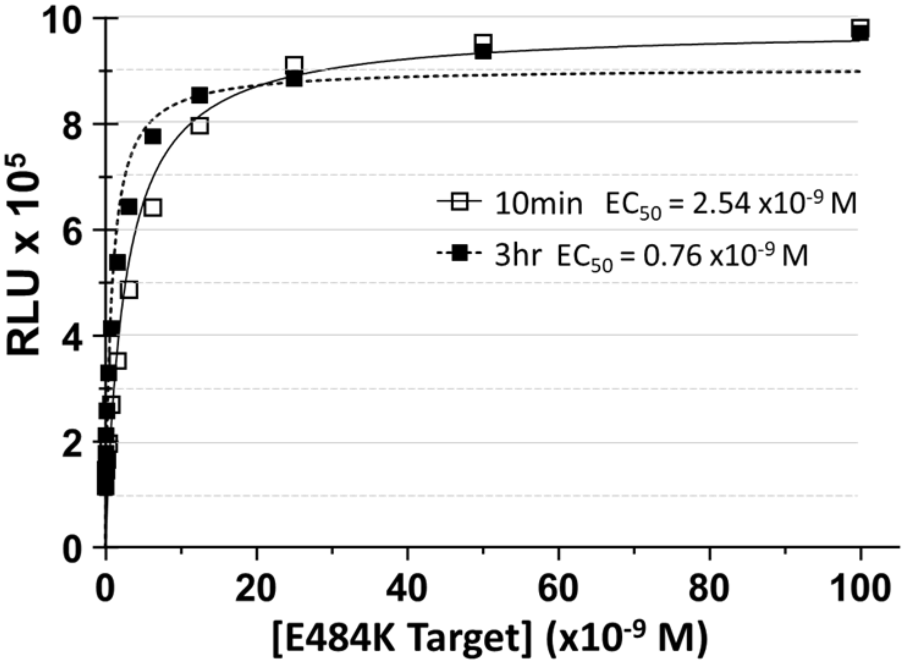


**Supporting Figure 3.** Bioluminescence of Eb.19 (E484K) with increasing target oligo concentration after 10 min or 3 hr incubation. Data were fit to a hyperbolic binding isotherm to calculate $EC_{50}$ values. The $EC_{50}$ was $2.54\times10^{-9}$ M for 10 min incubation; for 3 hr, $EC_{50}$ was $0.11\times10^{-9}$ M. Eb.19 (E484K) was present at $8\times10^{-11}$ M.

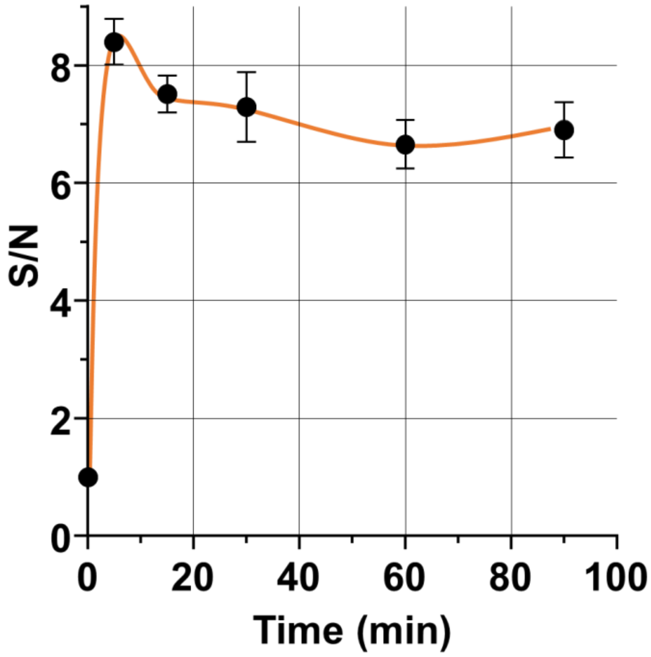


**Supporting Figure 4.** Unquenching of Eb.2' by digestion of the hairpin component of Eb.2' with DNase-1 produced similar S/N ratios compared to hybridization driven unquenching. Eb.2' ($1\times10^{-8}$ M, final) was mixed with DNase 1 (6 units) and DNase 1 buffer (from New England Biolabs)at 25 °C and analyzed for bioluminescence at selected intervals. The S/N was calculated from the ratio of bioluminescence in samples +/- DNase 1. The incubation period varies from 0 min to 90 min.

### 1. Materials:

#### Chemical

All chemicals were obtained from commercial suppliers and used directly unless otherwise mentioned: Fos-Choline-12 (Anatrace); Imidazole (Acros Organics); Triethylammonium acetate (Calbiochem); Dimethyl sulfoxide (EMD Millipore Corp.); Acetonitrile, glycerol tris(2-carboxyethyl)phosphine, Tris Base (Fisher Scientific Inc.); MeOH (Macron Fine Chemicals); Ampicillin, Isopropyl β-d-1-thiogalactopyranoside (MP Biomedicals); Nano-Glo® Luciferase Assay System (Promega); N-(3-Dimethylaminopropyl)-N′-ethylcarbodiimide hydrochloride, n-butanol, LB agar (miller), Luria Bertani Broth (Sigma); 23, 24-BISNOR-5-CHOLENIC ACID-3β-OL (Steraloids); KCl, $MgCl_2$ (VWR). The CPG solid supports (3'-dabcyl-CPG or dT-CPG, 1.0 µM), DNA phosphoramidites (dT-CE, dABZ-CE, dCAc-CE and dGdmf-CE), 5'-carboxy modifier-C10 amidite, 0.25 M tetrazole in acetonitrile, capping and oxidation solutions(Glen Research).

#### Plates

Corning® 96 Well Black Polystyrene Microplate （#3650）

Dialysis chambers: EMD Millipore Corp. D-Tube™ Dialyzer Maxi, MWCO 12-14 kDa (#71510)
Concentrators: Corning® Spin-X® UF 6 mL Centrifugal Concentrator, 5,000 MWCO Membrane. (#431482)

#### Buffers

Bacterial Cell Lysis buffer: 0.5 % Triton X-100, 0.05 M $K_2HPO_4$, 0.4 M NaCl, 0.1 M KCl, 10 % glycerol, 0.01 M imidazole, pH=7.3.

Ni-NTA Bind buffer: 1 M NaCl, 0.04 M $Na_2HPO_4$, 0.06 M imidazole, 20 % glycerol, pH=7.5.

Ni-NTA Elution buffer: 0.02 M $Na_2HPO_4$, 0.5 M NaCl, 0.5 M imidazole, 10% glycerol, pH=7.3.

Agarose gel extraction buffer: 20mM Tris HCl, pH=7.4.

Luciferase Assay DNA hybridization buffer: 100 mM KCl, 1mM $MgCl_2$, 10 mM Tris HCl, pH=8.0.

### 2. Methods:

#### 2A. Protein expression/ purification

Two C-terminal His-tagged Nluc-HhC precursor constructs were used in this study. The first construct Nluc-HhC has been described previously [1]. The second construct, Nluc-HhC(D46H)-SUMO, differs in using a gain-of-function HhC mutant [2] along with a SUMO tag for enhanced expression and solubility. E. coli BL21(DE3) containing expression plasmid for each His-tagged Nluc-HhC precursor was grown at 37 °C in 50 ml of LB broth with carbenicillin (100 μg/ml) and 250 RPM shaking. Once $OD_{600}$ reached 0.6-0.8, IPTG was added to the culture (0.5 mM, final) to induce expression. After 18-20 hours at 16°C, bacterial cells were harvested by centrifugation at 10,000 RPM for 10 mins and the pellet was resuspended in bacterial lysis buffer (3 ml). After 3 freeze (-80 °C) / thaw cycles and sonication, insoluble material was removed by centrifugation at 10,000 RPM for 60 min. To the clarified lysate, an equal volume of ice chilled 2x Ni-NTA bind buffer was added. The solution was applied to Ni-NTA spin columns (GE Health). His-tagged precursor was purified according to the manufacture's protocol. Purified Nluc-HhC or Nluc-HhC(D46H)-SUMO precursor protein was stored at -80 °C in Ni-NTA elution buffer with added TCEP ($5\times 10^{-3}$ M).

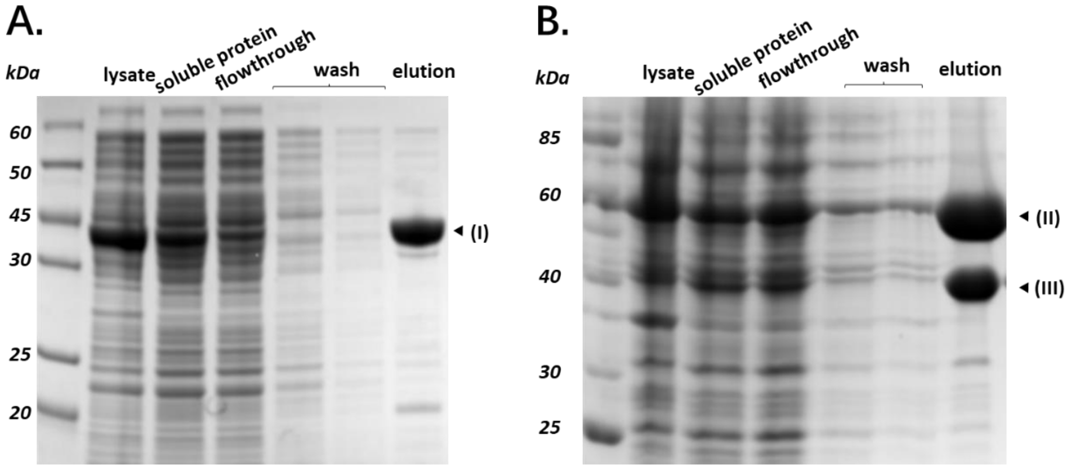


**Supporting Figure 5.** SDS-PAGE images showing the results of Nluc-HhC precursor protein purification. (A) Nluc-HhC precursor protein purification. (I) Nluc-HhC (44kDa). (B) Nluc-D46H-SUMO precursor protein purification. (II) Nluc-HhC(D46H)-SUMO (58kDa), (III) D46H-SUMO (39kDa). The presence of (III) is caused by spontaneous self-cleavage (hydrolysis) of the autoprocessing domain, HhC(D46H)-SUMO, fragment from Nluc.

### 2B. Solution-based EDC oligonucleotide sterylation (used for EB.1-3)

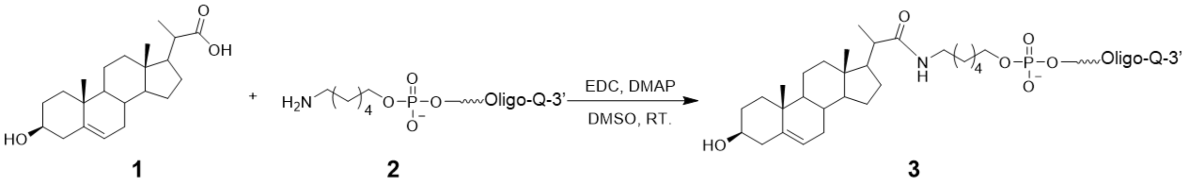


Q = Quencher

To 200 µl of DMSO, we first added 4.33 mg (12.5 µmol) of 23, 24-bisnor-5-cholenic acid-3β-ol **(1)** and 23.95 mg (125 µmol) of N-(3-Dimethylaminopropyl)-N′-ethylcarbodiimide hydrochloride (EDC). After incubation at room temperature for 30 min, 1.5 mg (12.5 µmol) of 4-Dimethylaminopyridine (DMAP) was added and incubated at room temperature for another 5 min. Last, 50 µl of 100 µM 5'-amino modified oligo **(2)** (in $H_2O$) was added and the solution was vortexed gently at room temperature overnight. The coupling reaction was extracted by adding 20 µl of 3 M sodium acetate, pH 5.2, and 1 ml of n-butanol. After a brief vortex and incubation at -80 °C for 1 hour, oligonucleotide was collected as a precipitate by centrifugation at 14,000 RPM for 20 min. The pellet was resuspended in 50 µl of water. Sterylated oligonucleotide (steramer) **(3)** was separated from sterol-free oligonucleotide by RP-HPLC, see below.

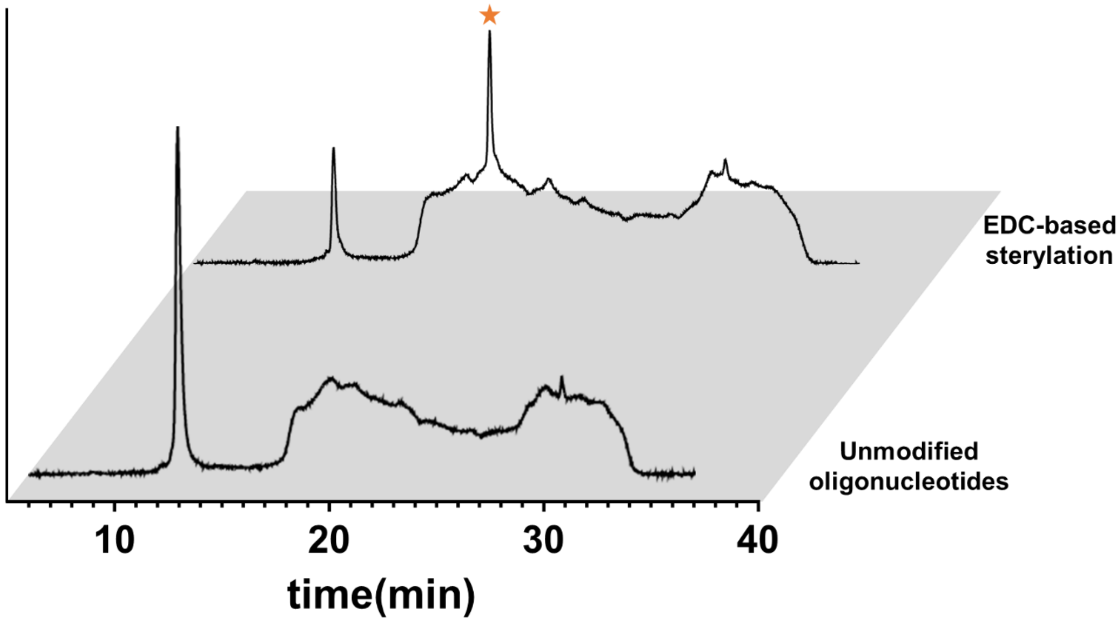


**Supporting Figure 6.** Reversed phase HPLC chromatogram of the sterol modified oligonucleotide with 3' quencher prepared by EDC-based sterylation**.** Sterol modified oligonucleotides (starred).

**2C. Synthesis and purification of sterol amine (5) for solid phase oligo coupling (for EB.19s)**

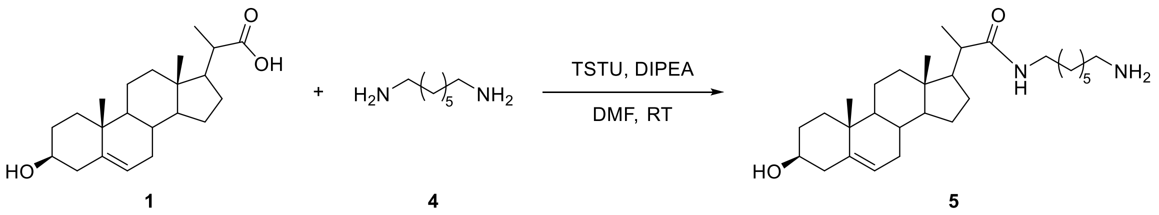


To 6 ml of anhydrous DMF, 173.5 mg (0.5 mmol) of 23, 24-bisnor-5-cholenic acid-3β-ol **(1)**, 435 ul (2.5 mmol) of N,N-Diisopropylethylamine (DIPEA), and 165.6 mg (0.55 mmol) of N,N,N′,N′-Tetramethyl-O- (N-succinimidyl) uronium tetrafluoroborate (TSTU) were added. The mixture was stirred at room temperature for 30 minutes. After that the mixture was added to 2 ml of 0.75 M 1,7-Diaminoheptane **(4)** in anhydrous DMF dropwise. Then the final reaction mixture was stirred at room temperature for 20 hours. After that the reaction mixture was washed with dichloromethane (10 ml) and saturated $NaHCO_3$ solution (10 ml) for three times, dried over $Na_2SO_4$, and evaporated under $N_2$ flow to afford 167.7 mg crude sterol amine **(5)** as a white to yellow solid.
The crude sterol amine **(5)** was dissolved in 1 ml of methanol and purified over a Restek viva C18 5 μm HPLC column (250×4.6 mm) using a gradient elution from 0% acetonitrile to 100% acetonitrile over 25 minutes. The flow rate was 1 mL/min and eluate was monitored at 210 nm. Sample injection volume was 100 μl. After HPLC purification, the solvent was evaporated to afford sterol amine **(5)** (144.6 mg, 63%) as a white solid.

**NMR Characterization of (5):**
$^{1}$H NMR spectra were acquired with Bruker Avance III HD 400 (400MHz) spectrometer at 25 °C. $CD_3OD$ (Sigma) was used as NMR solvent. 1H chemical shifts are reported as δ in units of parts per million (ppm) relative to methanol-d (3.31, s)
1H NMR (400MHz, $CD_3OD$) δ 5.34 (dd, 1H), 3.40 (m, 1H), 3.16 (m, 1H), 3.11 (m, 1H), 2.91 (t, 2H), 2.17 (dd, 1H), 1.38 (s, 6H), 1.14 (d, 2H), 1.03 (s, 3H), 0.75 (s, 3H). 13C NMR (100MHz, $CD_3OD$) δ 178.21, 140.86, 120.93, 71.00, 56.47, 52.61, 50.30, 43.64, 42.02, 41.60, 39.60, 39.30, 38.53, 37.14, 36.29, 31.88, 31.58, 30.89, 28.87, 28.34, 27.13, 26.98, 26.27, 25.96, 23.94, 20.76, 18.48, 16.56, 11.09.

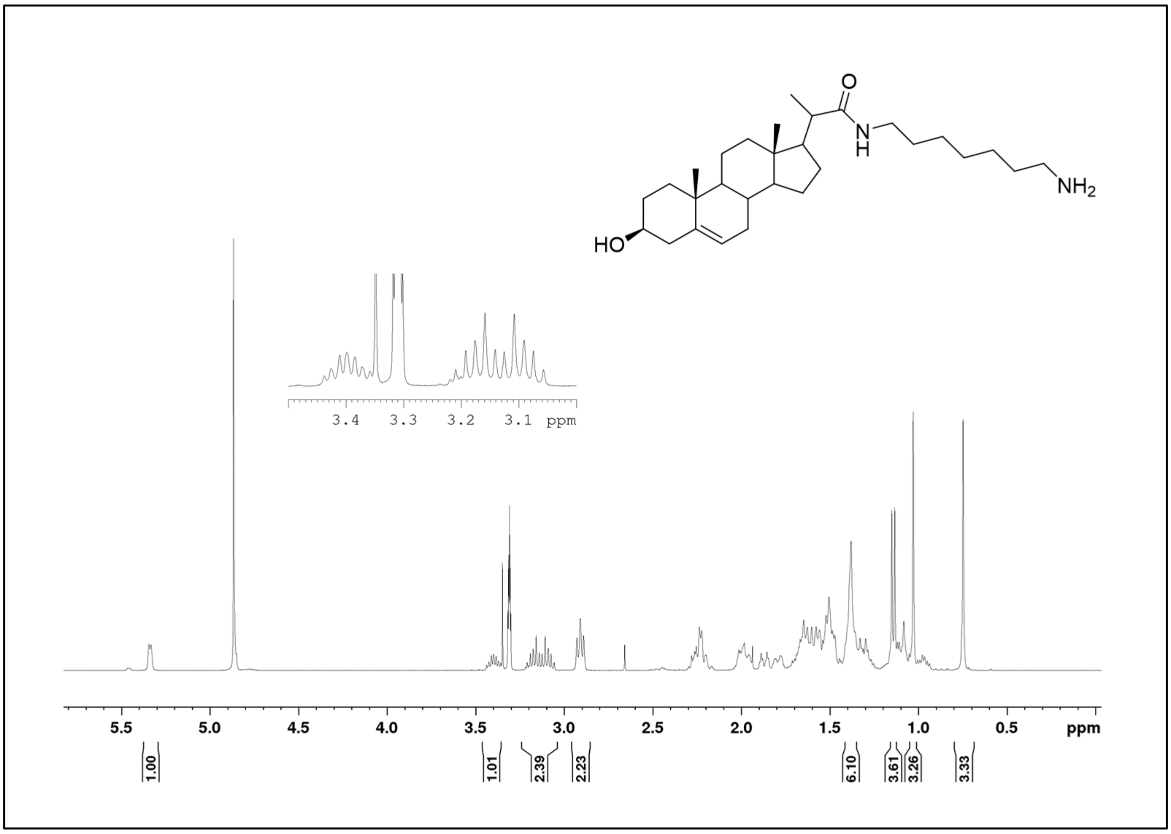


**Supporting Figure 7.** $^{1}$H NMR of the synthetic sterol amine (**5**).

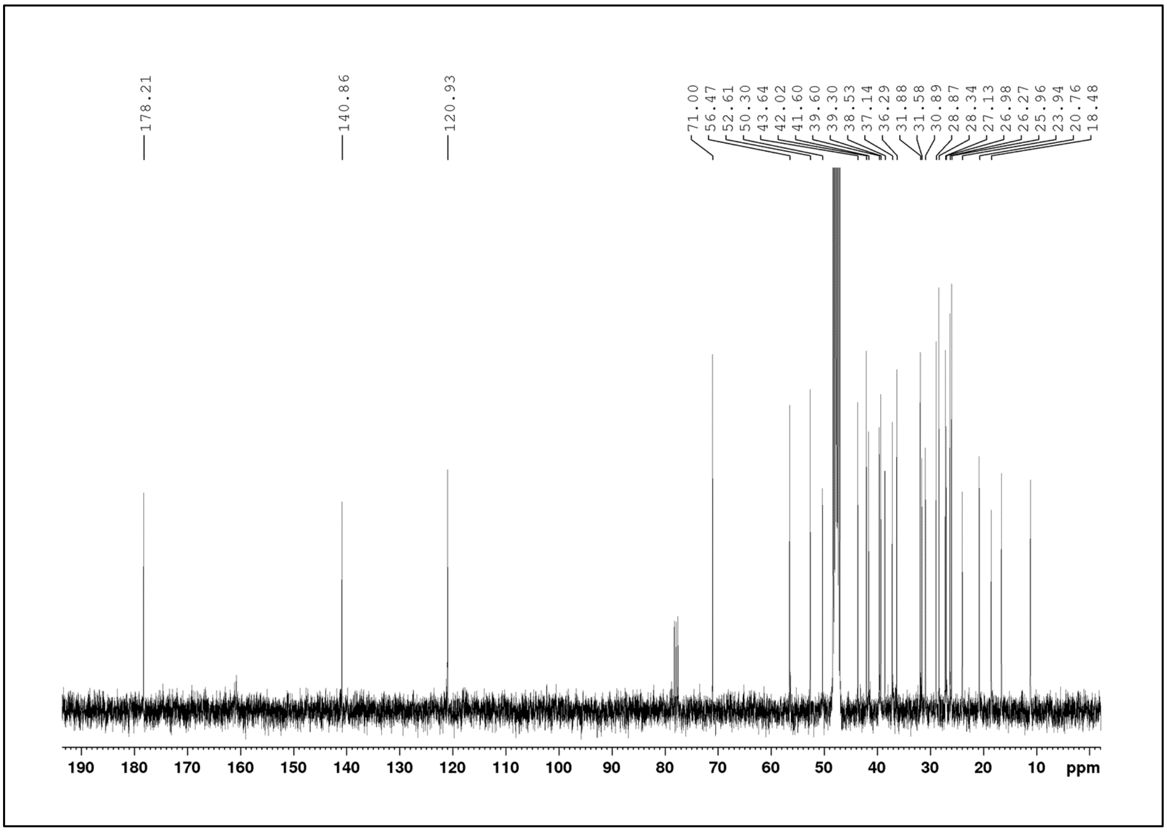


**Supporting Figure 8.** $^{13}$C NMR of the synthetic sterol amine (**5**)

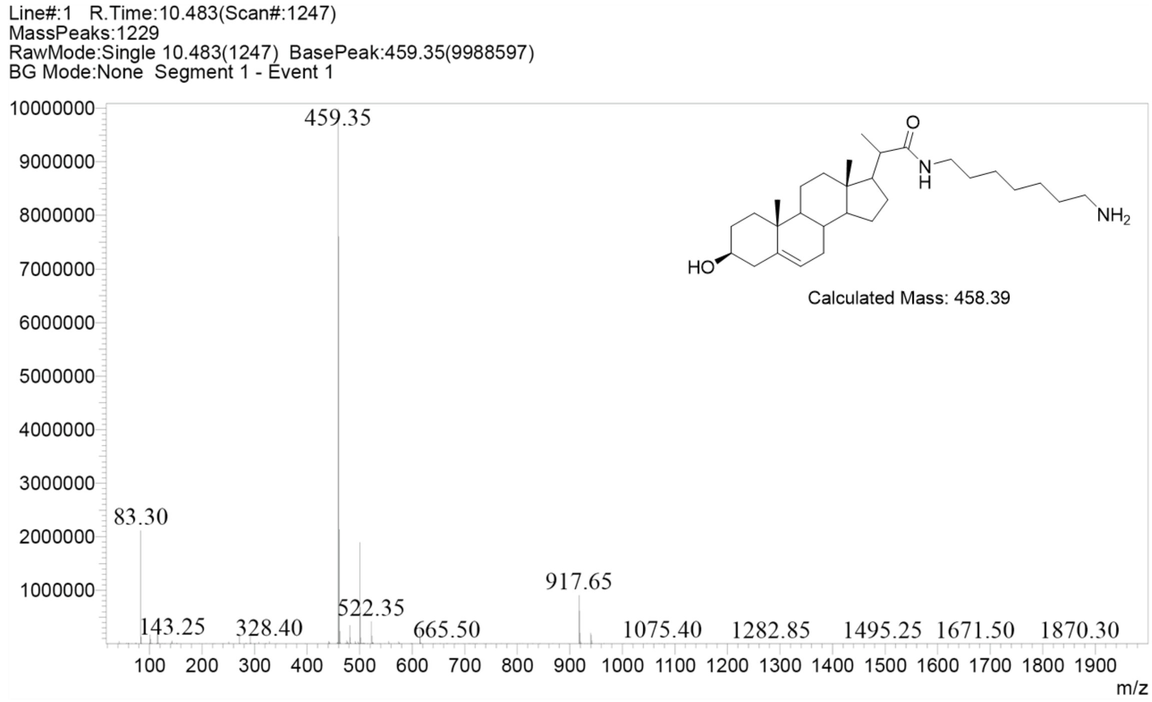


**Supporting Figure 9.** LC-MS analysis of the synthetic sterol amine (**5**).

### 2D. Solid Phase Steramer Synthesis

The sterylated-DNA oligomers (steramers, Table 1) were synthesized on a 1.0 µM scale (3'-dabcyl-CPG or dT-CPG) using standard DNA synthesis protocols on an automated Expedite 8909 DNA/RNA synthesizer. After synthesizing the desired DNA sequence, the 5'-end was modified with 5'-caroboxy modifier-C10 carrying a reactive NHS ester. The final detritylation step was eliminated after coupling with the 5'-carboxy modifier-C10. Prior to the next coupling with sterol amine, the solid support was washed with acetonitrile followed by drying with nitrogen purging for 5 minutes.

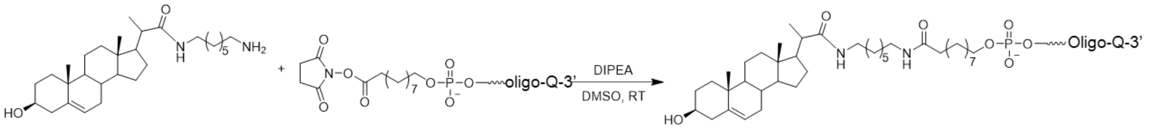


5 6 7

Q = Quencher

The manual coupling of sterol amine **(5)** (10 eq.) with the DNA-NHS ester on the CPG solid support was performed in anhydrous DMSO containing 10% DIPEA at room temperature for 2 hours using the push-pull syringe method. After the coupling, the column was thoroughly washed with 5×1 mL of acetonitrile and dried.
The steramers **(7)** were cleaved from the solid support using 40% methyl amine in water (2×0.8 mL) at room temperature for 2 hours. The cleavage solution was lyophilized to get the crude steramers. The exposure to light was avoided as much as possible while handling

the 3'-dabcyl labeled steramers synthesis. The identity of all the synthesized crude steramers were confirmed by the MALDI-ToF mass analysis (Table 1). The crude samples of all the steramers showed ~85-90% purity by RP-HPLC.

**Supporting Table 1: List of steramers prepared by solid phase synthesis**

| E-beacons | *Sequence (5'-3') | Mass calcd./found |
|---|---|---|
| Eb.2' | *sterol*- CGCTC CCAAAAAAAAAAACC GAGCG- dabcyl | 8780/8783 |
| Eb.19 (E484) | *sterol*- CGCTC TGGTGTT**GAA**GGTTT GAGCG- dabcyl | 8909/8912 |
| Eb.19 (E484K) | *sterol*- CGCTC TGGTGTT**AAA**GGTTT GAGCG- dabcyl | 8893/8898 |

* The underlined regions self-anneal to form the hairpin stem. The nucleotides in black color were for the target RNA/DNA detection. Sterol moiety at the 5'-end was introduced for the bioconjugation with nanoluciferase and the dabcyl unit at 3'-end was as a dark fluorescence quencher.

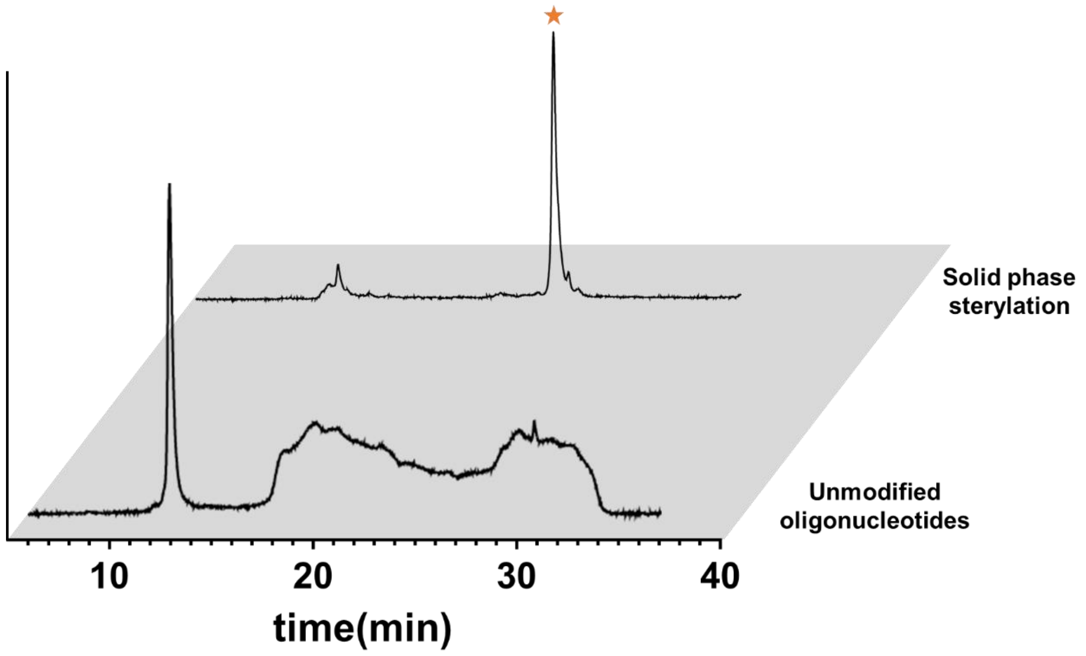


**Supporting Figure 10.** Reversed phase HPLC chromatogram of the sterol modified oligonucleotide with 3' quencher prepared by solid phase.

### 2E. General Conditions for E-beacon bioconjugation by HhC

To prepare Nluc-hairpin nucleic acid conjugates, Nluc-HhC precursor protein ($2\times10^{-6}$ M, final), Fos-choline 12 ($1.5\times10^{-3}$ M, final), Tris(2-carboxyethyl) phosphine hydrochloride (TCEP, $5\times10^{-3}$ M, final), Bis-Tris buffer (0.05 M, final, pH 7.1), ethylenediaminetetraacetic acid (EDTA, $5\times10^{-4}$ M, final), NaCl (0.1 M, final) were mixed. To that solution, Steramer ($1\times10^{-4}$ M, final) was added and the reaction was incubated at 16 °C overnight.

### 2F. E-beacon isolation by agarose gel extraction

Agarose gel extraction was used as a rapid means of isolating E-beacon. E-beacon conjugation reaction (100 µl) was combined with 6X gel loading buffer (20 µl) and separated on 2% agarose gel containing GelRed® Nucleic Acid Gel Stain (followed the manufacturer's "precast protocol"). The gel was run at 90 V in 1 x TAE buffer until the sample loading dye front was approximately 95% toward the end of the gel. E-beacon conjugate was visualized with BioRad Gel Doc® (UV tray), then excised, diced, transferred to an Eppendorf tube and soaked in Tris buffer (3 ml, 20 mM pH7.4) at 4 °C overnight. After centrifugation to gel fragment, the supernatant was concentrated by a Spin-X® UF Concentrator (Corning), 5 kDa MWCO.

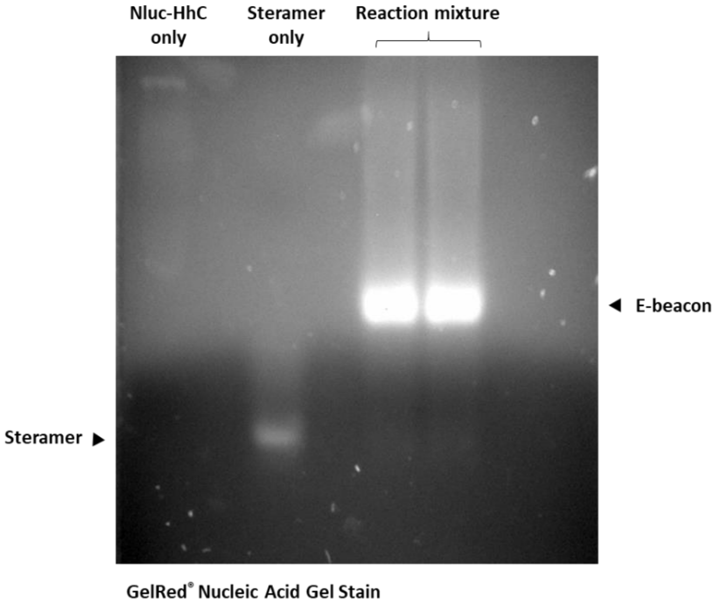


**Supporting Figure 11.** Image of the agarose gel for E-beacon purification. Nucleic acid moieties in the agarose gel were visualized by GelRed® nucleic acid gel stain giving bright bands.

### 2G. E-beacon Bioluminescence Measurements

To perform luminescence reading, 25 µl of E-Beacon solution was combined with 25 µl of sample nucleic acid in the DNA hybridization buffer a Corning® 96 Well Black Polystyrene Microplate. After incubation at 25 °C for the selected interval, we added 25 µl of Nano-Glo® Luciferase Assay System substrate (Promega). After another 5 min, bioluminescence was measured using a Synergy H1 Hybrid Multi-Mode Microplate Reader (BioTek).

### References

1. Zhang, X.; Xu, Z.; Moumin, D. S.; Ciulla, D. A.; Owen, T. S.; Mancusi, R. A.; Giner, J. L.; Wang, C.; Callahan, B. P., Protein-Nucleic Acid Conjugation with Sterol Linkers Using Hedgehog Autoprocessing. *Bioconjug Chem* **2019,** *30* (11), 2799-2804.

2. Zhao, J.; Ciulla, D. A.; Xie, J.; Wagner, A. G.; Castillo, D. A.; Zwarycz, A. S.; Lin, Z.; Beadle, S.; Giner, J. L.; Li, Z.; Li, H.; Banavali, N.; Callahan, B. P.; Wang, C., General Base Swap Preserves Activity and Expands Substrate Tolerance in Hedgehog Autoprocessing. *J Am Chem Soc* **2019,** *141* (46), 18380-18384.
